# Event Detection and Classification from Multimodal Time Series with Application to Neural Data

**DOI:** 10.1101/2023.12.20.572485

**Authors:** Nitin Sadras, Bijan Pesaran, Maryam M. Shanechi

**Affiliations:** Ming Hsieh Department of Electrical and Computer Engineering, Viterbi School of Engineering, University of Southern California, Los Angeles, CA, USA; Departments of Neurosurgery, Neuroscience, and Bioengineering, University of Pennsylvania, Philadelphia, PA, USA; Neuroscience Graduate Program, University of Southern California, Los Angeles, CA, USA

**Keywords:** multimodal, spiking activity, local field potentials (LFP), neural decoding, maximum likelihood

## Abstract

The detection of events in time-series data is a common signal-processing problem. When the data can be modeled as a known template signal with an unknown delay in Gaussian noise, detection of the template signal can be done with a traditional matched filter. However, in many applications, the event of interest is represented in multimodal data consisting of both Gaussian and point-process time series. Neuroscience experiments, for example, can simultaneously record multimodal neural signals such as local field potentials (LFPs), which can be modeled as Gaussian, and neuronal spikes, which can be modeled as point processes. Currently, no method exists for event detection from such multimodal data, and as such our objective in this work is to develop a method to meet this need. Here we address this challenge by developing the multimodal event detector (MED) algorithm which simultaneously estimates event times and classes. To do this, we write a multimodal likelihood function for Gaussian and point-process observations and derive the associated maximum likelihood estimator of simultaneous event times and classes. We additionally introduce a cross-modal scaling parameter to account for model mismatch in real datasets. We validate this method in extensive simulations as well as in a neural spike-LFP dataset recorded during an eye-movement task, where the events of interest are eye movements with unknown times and directions. We show that the MED can successfully detect eye movement onset and classify eye movement direction. Further, the MED successfully combines information across data modalities, with multimodal performance exceeding unimodal performance. This method can facilitate applications such as the discovery of latent events in multimodal neural population activity and the development of brain-computer interfaces for naturalistic settings without constrained tasks or prior knowledge of event times.

## 1. Introduction

Event detection from time-series data is an important and well-studied signal-processing problem. When we observe time series data that contain a known event-related template signal with an unknown delay, the problem of event detection is to estimate this delay or event time. When the observed time series can be modeled as this known template signal in Gaussian noise, a traditional matched filter can be used to perform event detection [1; 2]. In various applications, however, the observed time series are multimodal, consisting of both Gaussian and point-process signals. In neuroscience experiments, for example, collected datasets can consist of simultaneous local field potentials (LFPs) and neuronal spiking activity, which can be modelled as Gaussian and point-process signals, respectively. While methods have been developed for event detection from each of these single modalities in isolation [2–5], no such method exists for multimodal time series containing a mixture of Gaussian and point-process signals.

A specific neuroscientific application of interest is that of brain-computer interfaces (BCIs), which are systems that aim to decode motor or cognitive states from a user’s neural activity to restore or augment task performance [6–25]. Prior work has shown that motor and cognitive states are represented across multiple spatiotemporal modalities of neural activity, and that BCI decoding of these states can significantly benefit from using multimodal data [26–45]. Such BCI decoders are often trained and tested either on stereotyped tasks or on trial-based data that is aligned, or ‘time-locked’, to a known event such as the onset of a visual stimulus during a cognitive task. In real-world applications, however, it is likely that this event time is unknown and time-locking therefore cannot be performed. In order to enable high-performance multimodal motor or cognitive BCIs when event times are unknown or tasks are not stereotyped, a method to simultaneously detect and decode events from multimodal signals is necessary.

The problem of event detection from multimodal time series is challenging because of the differences in statistical properties of Gaussian and point-process signals [31; 46; 47] and because the events whose times are unknown can also have different yet unknown classes. In order to address this challenge, we develop the multimodal event detector (MED), an algorithm which performs event detection in multimodal time series with multiple unknown event classes. We do this by deriving the maximum-likelihood estimator of simultaneous event times and classes from multimodal data. We first write a parametric likelihood model of Gaussian and point-process time series that encode an event with an unknown time and class, learn the model parameters from data, and then derive the estimate of the time and class that maximize this likelihood function. We validate this method in simulations and in spike-LFP neural datasets recorded from a nonhuman primate (NHP) performing a rapid eye-movement (saccade) task, with the goal of detecting the times and directions of saccades. In simulated and real data, we show that the MED can successfully detect and classify eye movements from multimodal spike-field data. We further show that the MED successfully integrates information from both data modalities, with performance increasing as signal channels of either modality are added.

## 2. Methods

We first describe our model of how event times and classes are encoded in multimodal time series. We then derive the MED as the maximum-likelihood estimator of both event times and classes.

### 2.1. Multimodal Model

Point process signals can be described as a time-series of 0’s and 1’s. These point processes can be characterized by a conditional intensity function (CIF) that models the rate *λ* at which nonzero values, or ‘spikes’, occur as a function of external covariates. We model point processes via a Poisson generalized linear model (GLM), where the logarithm of the CIF is a linear function of the covariates. Poisson point-process models have been successfully used to model neuronal spiking activity [48–59]. Further, prior work has developed Poisson GLMs that explicitly encode event times and classes, and showed that such models are a good fit to the spiking of neurons that encode stimulus-related events [60]. However, such models have not yet been used to develop multimodal detection and decoding algorithms, as we do here.

In the context of event detection, the covariates that we model are the event time *t*_*s*_ ∈ [0, *T*] and the event class *s*, where [0, *T*] is the interval on which the event can occur and *s* is a ‘one-hot’ vector in ℝ^*S*^, where *S* is the number of possible classes. In a manner similar to [60], we write the point process CIF as

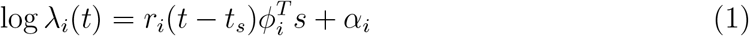

Here, *ϕ*_*i*_ ∈ ℝ^*S*^ and *r*_*i*_(*t*) are, respectively, the spatial and temporal responses of channel *i*, while *α*_*i*_ characterizes the baseline rate. The interpretation of this CIF is that an event at time *t*_*s*_ will evoke a temporal response *r*_*i*_(*t*) that is modulated by a spatial response 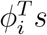. Given the CIF *λ*_*i*_(*t*) and assuming inhomogeneous Poisson statistics, the likelihood of observing a binary signal at channel *i* with spikes at times {*t*_*i,m*_}_*m*=1:*Mi*_ = {*t*_*i*,0_, …, *t*_*i,Mi*_} is given by

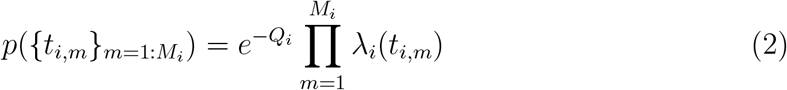

where 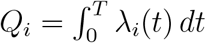, *m* is the spike time index, and *M*_i_ is the total number of spikes observed from channel *i* [61].

Similarly to our point process model, we model continuous channels as having spatial and temporal responses to an event at time *t*_*s*_ of class *s*:

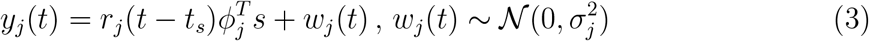

where *w*_*j*_(*t*) is Gaussian noise and 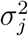 is the variance of this noise for channel *j*. The corresponding likelihood is

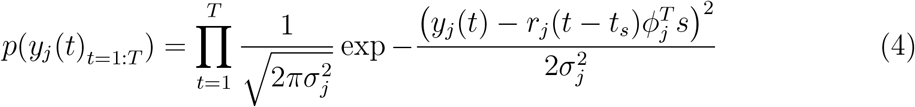

In order to form a joint model of Gaussian and Poisson channels, we assume that they are conditionally independent, given the stimulus time *t*_*s*_ and class *s*. With this assumption, the joint likelihood for *I* Poisson channels and *J* Gaussian channels is

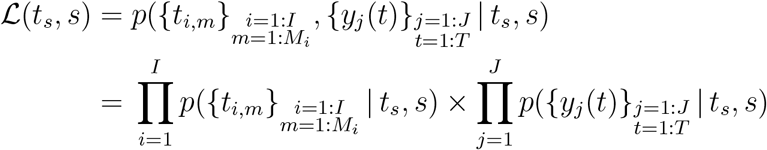

Substituting in our likelihood models from (2) and (4), we get

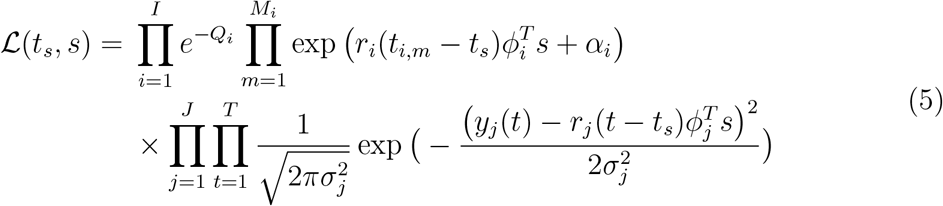

This provides a multimodal model of Gaussian and point-process signals that encode both event times, via a temporal response, and event classes, via a spatial response. We note that prior work has been done to model spatiotemporal responses for point process signals [5; 60], but the above does so for multimodal signals to solve the unaddressed problem of simultaneous event detection and classification from multimodal signals, as we do next.

### 2.2. Maximum Likelihood Estimate of Event Times and Classes

Using the models defined in section 2.1, we can now formalize the problem of simultaneous event detection and classification as an optimization problem. Specifically, our goal is to find the event time and class that maximize the log-likelihood of the observed multimodal data:

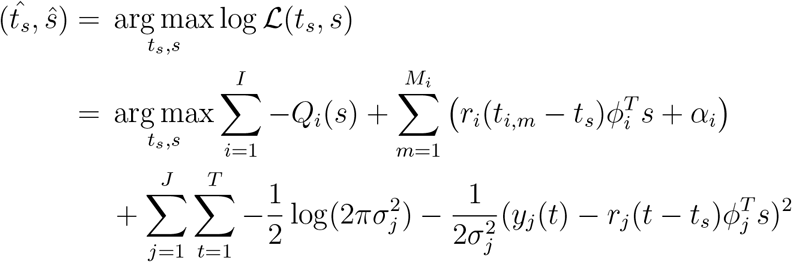

Removing additive terms that do not vary with *t*_*s*_ or *s*, and therefore do not impact the argmax, this simplifies to:

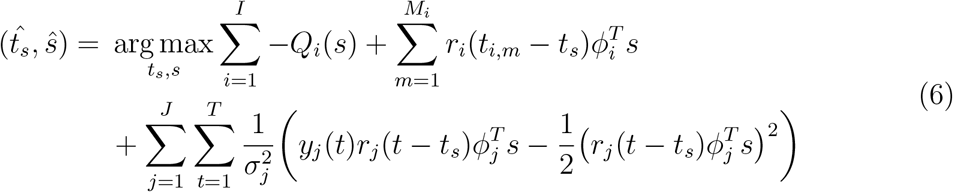

To further simplify this expression, we define *u*_*i*_(*t*) as the Poisson binary time series:

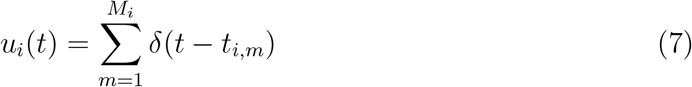

where *δ*(*t*) is the Dirac delta function. We can now re-write the following terms from (6) as convolutions:

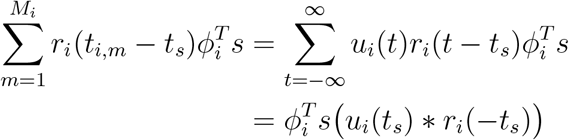

And

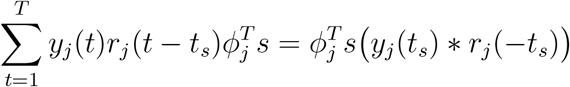

where ∗ represents convolution. Substituting these convolution expressions into (6), we get

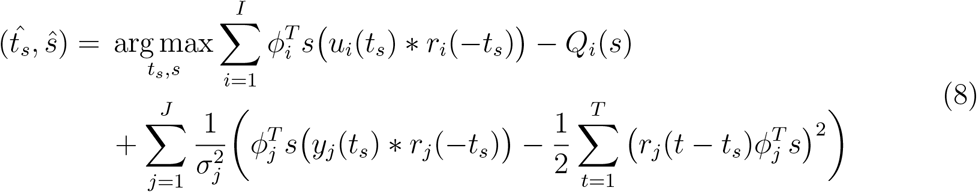

Finally, we note that if the support of *r*(*t*) is much smaller than *T*, then the term 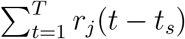 does not vary with *t*_*s*_. This is because delaying *r*_*j*_(*t*) by *t*_*s*_ samples does not change the value of its summation over the entire duration *T*, as long as the entire support of *r*_*j*_(*t*− *t*_*s*_) lies within [0, *T*]. Assuming that this is true, we can further simplify the second term in the sum over the *J* Gaussian channels in (8):

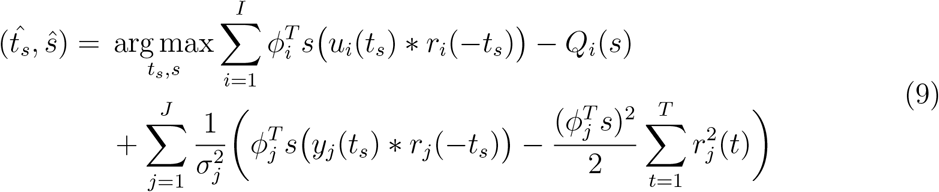

This is a sum of linear matched filters that are matched to the temporal responses *r*(*t*) and scaled by the spatial responses *ϕ*^*T*^ *s*. Point processing channels are vertically shifted by the term *Q*_*i*_(*s*), while Gaussian channels are vertically shifted by 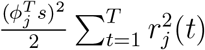 and then scaled by their inverse noise variance 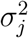. The shift terms set the baseline levels for each channel’s contribution to the MED output. The inverse noise variance scaling for Gaussian channels means that noisier channels have a smaller contribution to the MED output.

An additional challenge in multimodal integration when some modalities are discrete and some are continuous is that of proper scaling of likelihoods. In particular, while the point process likelihood for discrete random variables provides a probability measure (i.e. probability of a number of spikes), the Gaussian likelihood for continuous random variables provides a density measure that needs to be integrated over a range of values to provide a probability measure (in and of itself is not a probability). This distinction between discrete and continuous modalities makes the multimodal approach sensitive to scaling of the continuous signals (see details in section 4). To address this challenge and additionally account for model mismatch in real datasets, we introduce a single cross-modal scaling parameter *k* that weighs the contributions of the two data modalities in the likelihood model:

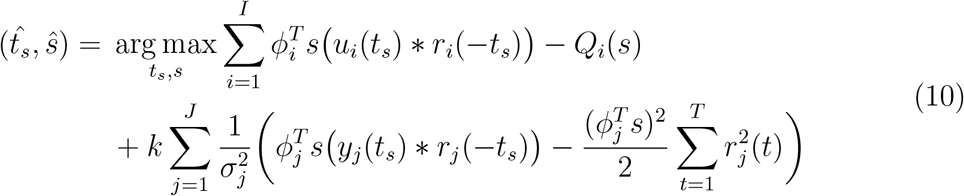

This scaling parameter can be learned via grid-search on training data by maximizing any performance metric of interest. Here, we choose the value of k that maximizes the sum of the event detection AUC (see section 2.5) and event classification accuracy on training data.

We can interpret the output of the MED as *S* separate signals, or one for each possible event class. At any given time, the maximum-valued output signal corresponds to the maximum-likelihood estimate of the event class, while the peaks in this maximum-valued signal correspond to the maximum-likelihood estimates of the event time. This idea is illustrated using simulated spike-field data in Figure 1, where correct saccade classification is indicated by the background color matching the color of the peak signal.

**Figure 1.**
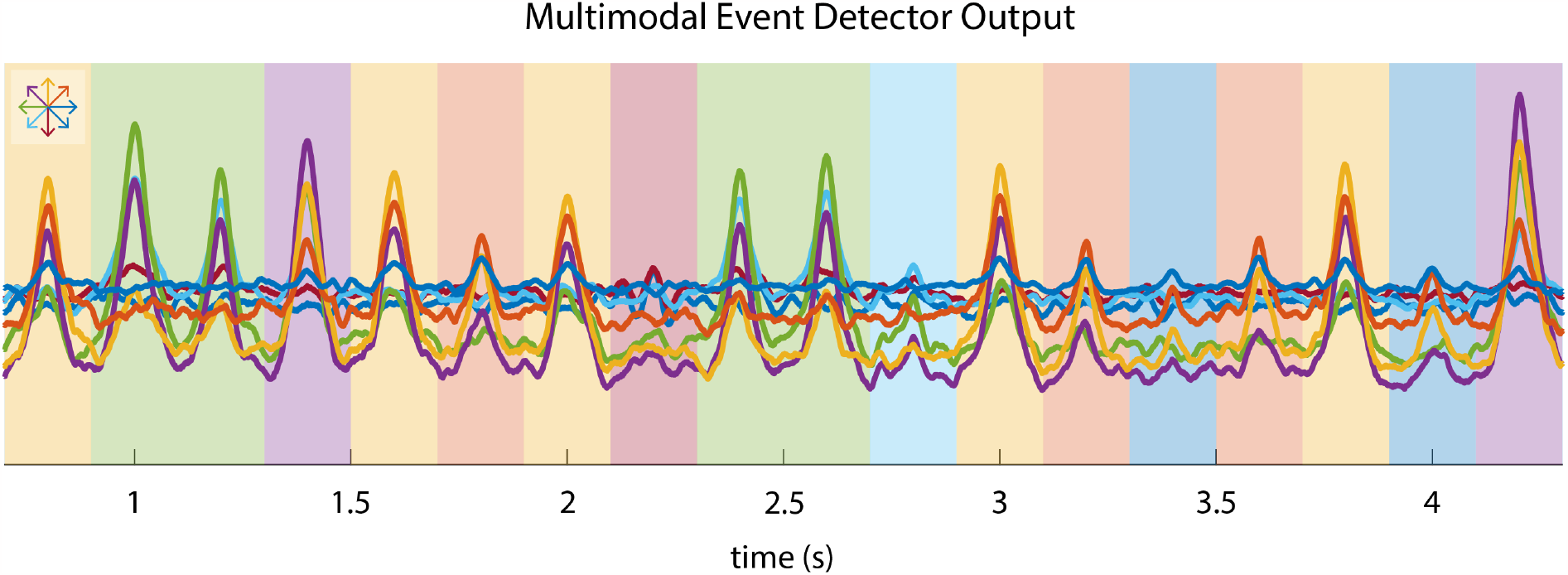
Sample output of the MED on simulated multimodal neural activity. In this example, the events are rapid eye movements (saccades), and the event classes are one of 8 eye movement directions. The MED produces 8 output signals, one for each class, which are shown plotted here. The color of the background indicates the true saccade direction at a given time, while the color of the maximum-valued output signal corresponds to the maximum-likelihood estimate of the saccade direction. Peaks in the maximum-valued signal correspond to the maximum-likelihood estimate of the saccade times. The mapping between colors and saccade directions is shown in the top left.

In order to use the MED in a real dataset, the parameters of our model in (5) must be estimated from training data. A maximum-likelihood method for estimating these parameters is described in Appendix A.

### 2.3. Multimodal Model of Saccade-Sensitive Neural Activity

Prior work has shown that saccadic eye movements elicit transient responses in neural activity that depend on the saccade direction [5; 60]. This provides an ideal validation setting for the MED by allowing us to test whether it can be used to detect and classify saccades from multimodal neural activity. In the dataset described in Section 2.4, there are 8 possible eye movement targets, and as such we consider the case where there are eight possible saccade directions. We model this by making our event class *s* a one-hot vector in ℝ^8^, with each entry representing one possible saccade direction. The spatial responses *ϕ* ∈ ℝ^8^ encode how saccades to each possible direction change the magnitude of the temporal response *r*(*t*). This saccade encoding model is illustrated in Figure 2A. Simulated multimodal data based on this model is shown in Figure 2B.

**Figure 2.**
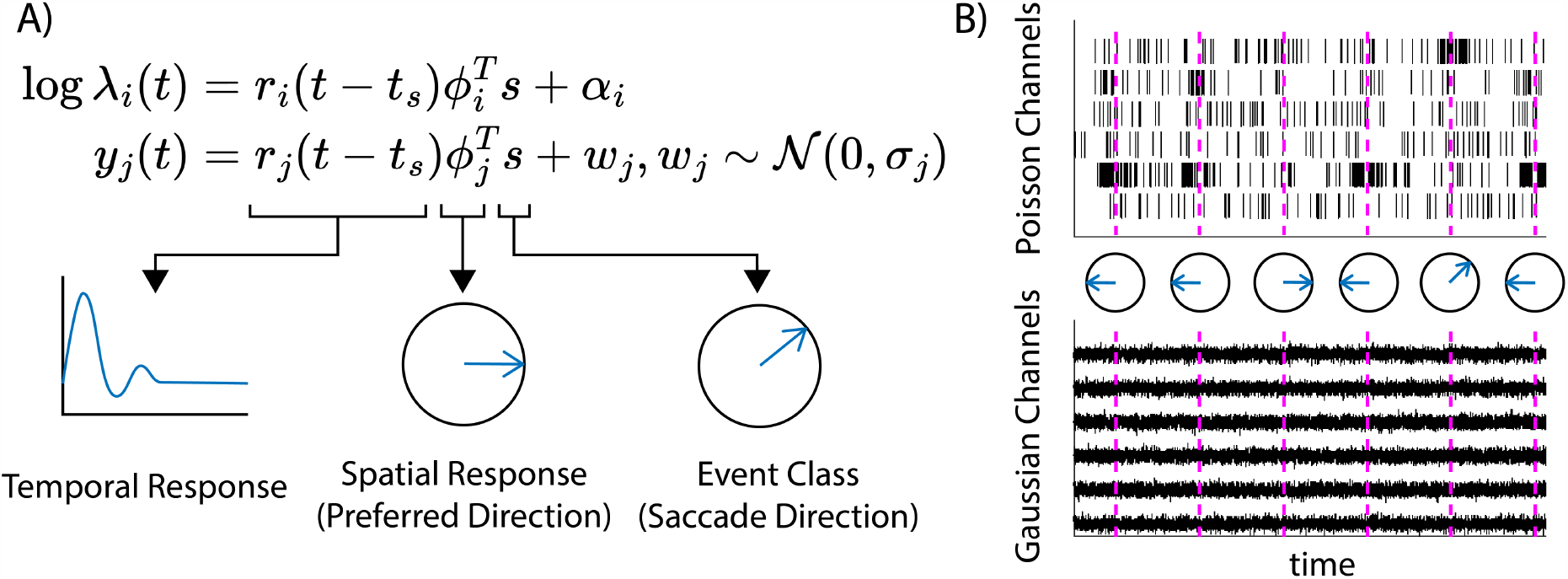
Model of saccade-sensitive multimodal neural activity. A) *r*(*t*) is the temporal response of the channel to an event at time *t*_*s*_, while the spatial parameter *ϕ* encodes the magnitude of the channel’s response to each possible saccade direction *s*. B) Simulated Poisson (top) and Gaussian (bottom) data based on the model in A. Saccade times are indicated by vertical dashed lines, and the corresponding saccade directions are indicated by the circled arrows.

### 2.4. Nonhuman Primate Saccade Task

In addition to simulations, we validated the MED on multimodal spike-LFP data collected from the prefrontal cortex (PFC) of an NHP performing a delayed saccade task. The task design is as follows. First, the NHP was required to maintain its gaze on a central point on a screen for 500-800ms. After this fixation, a target cue appeared in one of eight possible locations. After a delay period of 1000-1500ms, a ‘go’ cue prompted the NHP to make a saccade to the target. A movable electrode array consisting of 32 electrodes (Gray Matter Research, USA) was placed over the prearcuate gyrus of the lateral PFC to record neural activity. Raw neural signals were sampled at 30 kHz. In order to isolate single-unit activity, the raw data was preprocessed by highpass filtering at 300 Hz and thresholding at 3.5 standard deviations below the signal mean. Spike sorting was then performed via principal component analysis and k-means clustering. Eye position was recorded using an infrared eye-tracking system (ISCAN, USA) with a sampling rate of 120 Hz. All surgical and experimental procedures were in compliance with National Institute of Health Guide for Care and Use of Laboratory Animals and were approved by the New York University Institutional Animal Care and Use Committee. Further details can be found in [62].

### 2.5. Performance Evaluation

We assess the ability of the MED to detect event times using Receiver Operating Characteristic (ROC) curve analysis [63]. The ROC plots the probability of true detection of events against the probability of false detection as a threshold for detection is varied. The area under this curve (AUC) is used as a threshold-free performance metric, with an AUC of 0.5 indicating chance-level detection and an AUC of 1 indicating perfect detection. We use a modified version of the AUC that counts detections that are within 200ms of a true event to count as true detections.

The details of this modified AUC measure are as follows. Let *x*(*t*) be the maximum value of the MED’s output at any time *t*, and let *h* be an event detection threshold. We consider all peaks in *x*(*t*) that are above the threshold *h*. If a peak is within 200ms of an actual saccade, then it is considered a true positive. If it is not, then it is considered a false positive. In this way, we record the true positive rates (TPRs) and false positive rates (FPRs) as the threshold *h* varies from min(*x*(*t*)) to max(*x*(*t*)). We can then plot these FPR and TPR values against each other to construct an ROC curve, and the area under this curve is our AUC metric.

## 3. Results

We validated the MED in the context of saccade detection from neural activity, using both numerical simulations and nonhuman primate neural signals. In both cases, the goal of the MED is to detect saccade onset time and direction.

### 3.1. Simulation Results

We simulated the activity of five Poisson and five Gaussian channels using the model described in Section 2.3. We generated neural signals corresponding to 20 saccades, with a jittered 2 second gap between each saccade. The simulated point process channel signals had a minimum firing rate of 10 Hz and a maximum firing rate of 100 Hz, while Gaussian signals had an SNR of 0.1. These values were chosen in order to roughly match the properties of the real dataset we analyze in section 3.2. We repeated this simulation 104 times and for each repetition the true neural parameters, saccade directions, and saccade time jitters were randomized. The true neural parameters were used to generate training and test data for each iteration, and neural parameters were estimated from the training data using the method described in Appendix A. These estimated parameters were then used to perform simultaneous saccade time detection and direction classification on the test data.

A comparison of the estimated parameters with the ground truth is shown in Figure 3. The event detection and classification performance metrics, averaged across simulation iterations and as a function of multimodal channel counts, are shown in Figure 4. This simulation analysis shows that: 1) we can successfully estimate spatial and temporal response parameters from simulated multimodal data and 2) the MED can use these estimated parameters to successfully perform simultaneous event detection and classification from multimodal data. Importantly, the MED successfully combines information across data modalities, with performance increasing monotonically as channels of either modality are added.

**Figure 3.**
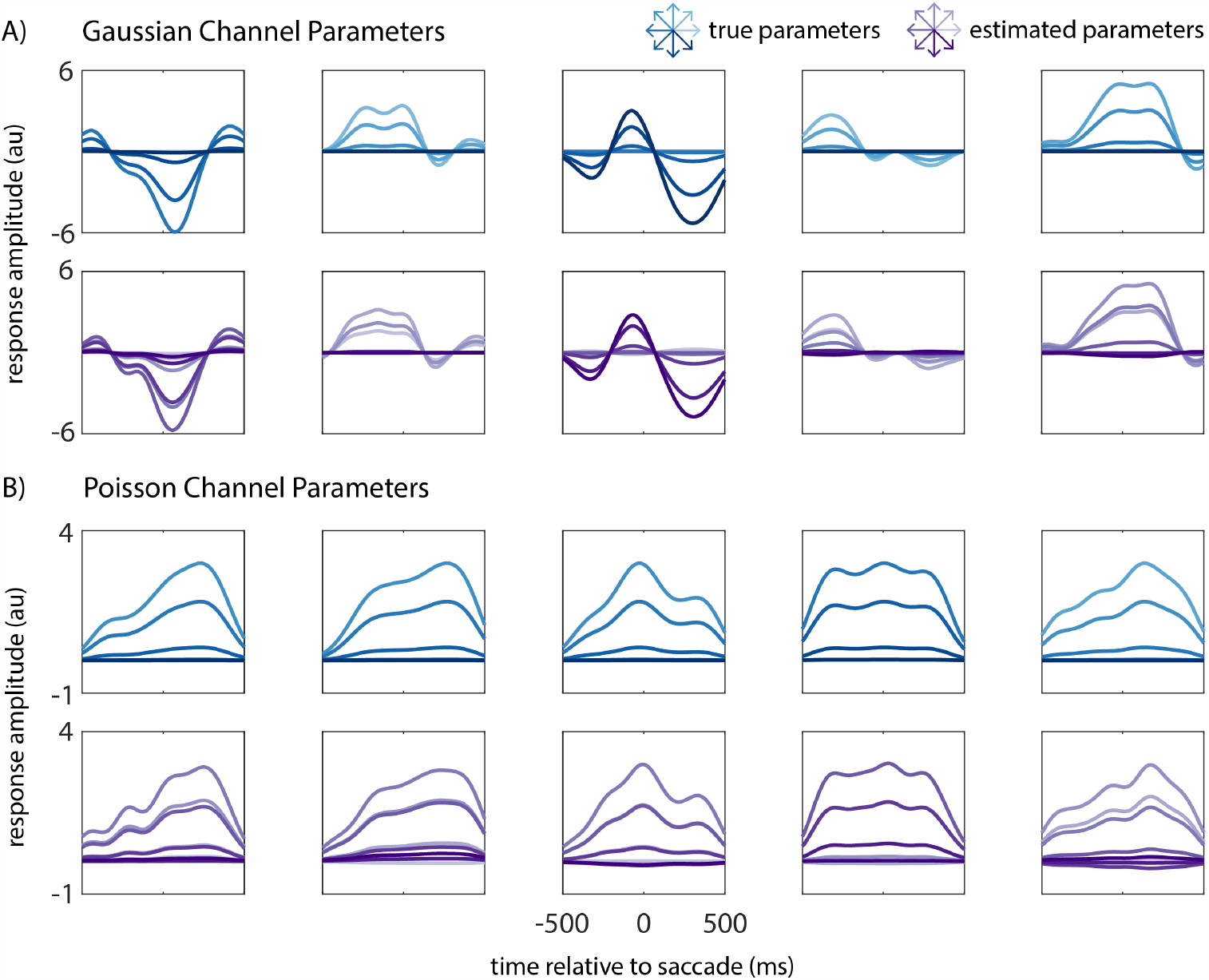
Comparison of true and estimated parameters based on simulated multimodal data. We simulated multimodal neural activity that encodes saccade time and direction, and used maximum likelihood estimation to estimate the model parameters. A) Ground truth (top row) and estimated (bottom row) spatiotemporal responses for five example Gaussian channels. Each trace indicates the channel’s response to a specific saccade direction, indicated by its color. The mapping between trace colors and saccade directions is shown on the top right. Our maximum likelihood estimation procedure resulted in parameters that closely match the ground truth. B) Same as A but for five example Poisson channels.

**Figure 4.**
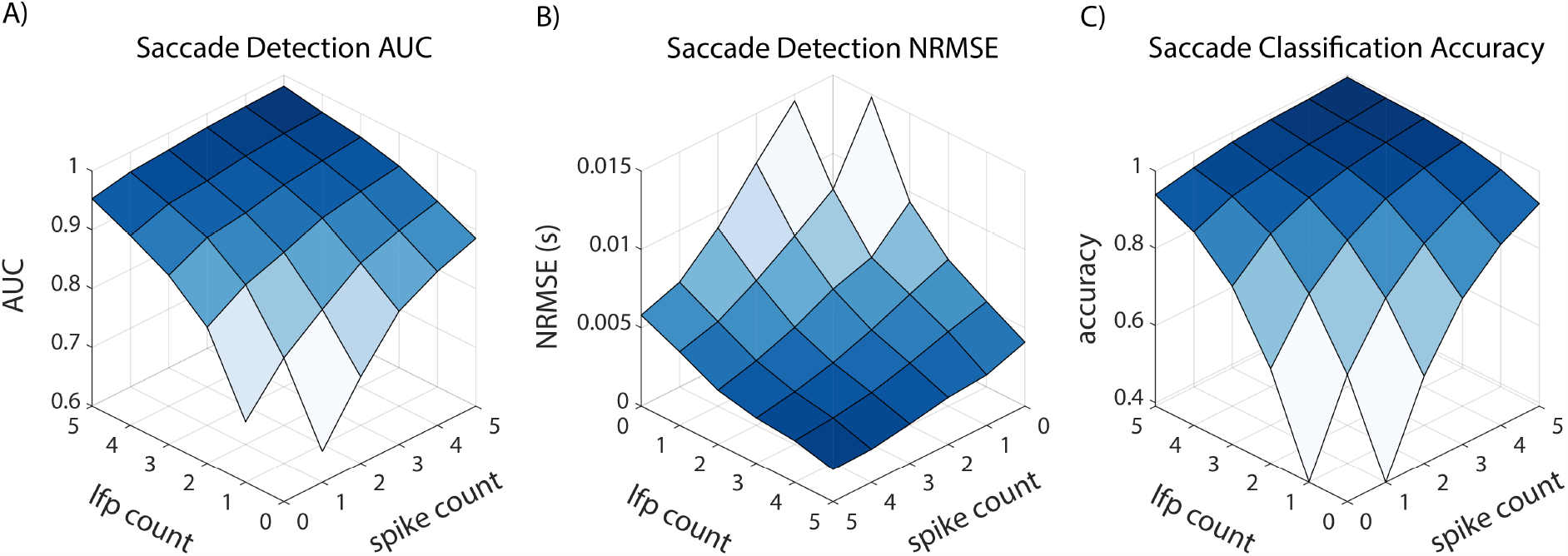
MED performance on simulated saccade-sensitive multimodal neural activity. All measures are shown as a function of the number of channels of each modality and are averaged over 104 simulation repetitions. A) saccade time detection area under the curve (AUC); higher AUC indicates better saccade detection performance. B) saccade time detection normalized root-mean-square error (NRMSE); lower NRMSE indicates better performance. C) saccade direction classification accuracy. The MED successfully combines information across data modalities, with performance increasing monotonically as channels of either type are added.

### 3.2. Nonhuman Primate Data Results

To validate the MED on a real-world dataset, we used spike-LFP neural data recorded from an NHP performing a delayed-saccade task. In this task, detailed in Section 2.4, the NHP first had to maintain its gaze on a central fixation point on a screen, and then make a saccade to one of eight peripheral targets after a ‘go’ cue [62]. We used a subset of this data consisting of eight continuous recording sessions containing a total of 1871 trials. We evaluated the MED via leave-one-out cross validation over sessions, where all sessions except for one were used to train the MED, and the remaining session was used to test it. This cross-validation was repeated eight times so that each session could be used for testing. In order to explore how the MED integrates information across modalities, we measured performance as channels of both modalities were added one at a time. For each cross-validation fold, we shuffled the order in which spike and LFP channels were added 10 times. These 10 shuffles were the same for each fold.

The MED was able to successfully detect and classify saccades simultaneously from multimodal spike-LFP data. First, even with a relatively low number (10) of spike and LFP channels, the MED achieved a saccade time detection AUC of 0.96 (chance level = 0.5) and a saccade direction classification accuracy of 0.55 (chance level = 0.125). These are both significantly higher (*p <* 5e-5, paired t-test, N=80) than the corresponding chance levels. Second, the MED successfully achieved multimodal fusion. Both detection AUC and classification accuracy increase as channels of either type are added, as shown in Figure 5A and B. Indeed, multimodal performance was significantly greater than both spike-only (p *<* 1e-3, Hochberg-corrected paired t-test, N=80) and LFP-only performance (p *<* 1e-14, Hochberg-corrected paired t-test, N=80) for all unimodal channel counts, as shown in Figure 5C and D. While this performance increase was indeed significant for all unimodal channel counts, the increase was larger in the low-information regime, i.e., when unimodal channel counts were smaller.

**Figure 5.**
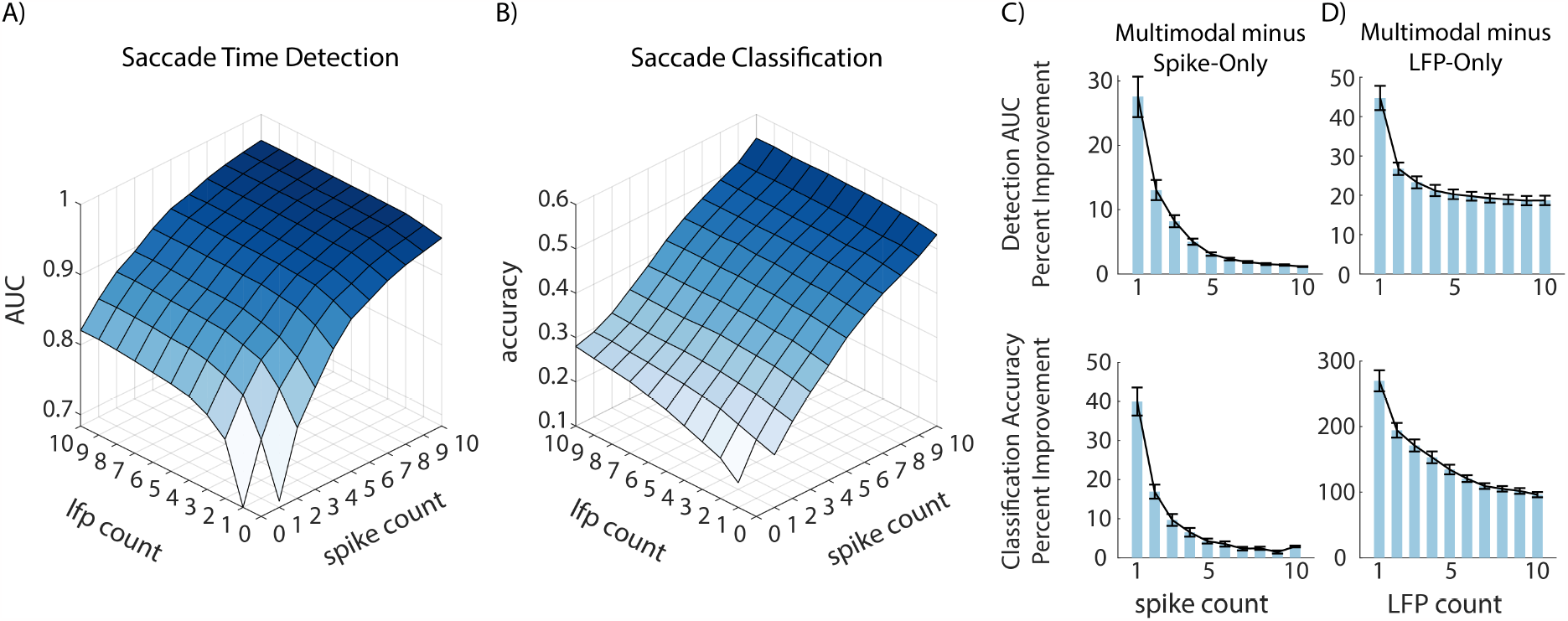
MED performance on multimodal nonhuman primate (NHP) neural data. A) Saccade time detection AUC. B) Saccade direction classification accuracy. Results in A and B are averaged over 8 cross-validation folds and 10 shuffles of channel order. C) Performance benefit of using multimodal data over spike-only data, as a function of the number of spike channels. Performance benefit is shown for both saccade time detection AUC (top) and direction classification accuracy (bottom). Bars indicate the mean percent improvement from spike-only to multimodal performance, and error bars indicate the standard error of the mean (SEM). Multimodal data includes all LFP channels, with different number of spike channels (i.e., spike count) as indicated on the x-axis. For all spike-count values, there was significant benefit in adding LFP channels (p *<* .001, paired t-test, N=80). D) Same as C, but showing the benefit of using multimodal data over LFP-only data. For all LFP-count values, there was significant benefit in adding spike channels (p *<* 1e-14, paired t-test, N=80).

Interestingly, despite LFP-only performance being lower than spike-only performance, the MED was still able to improve upon spike-only performance by adding LFPs. This indicates that the MED truly integrated information from both modalities, taking advantage of information present in LFP channels that was not present in spike channels. This result also suggests that spiking and LFP activities carry non-redundant information about both the timing and the class of a saccade event.

## 4. Discussion

In this work we developed the MED, which solves the unaddressed problem of event detection and classification from multimodal point-process and Gaussian time series. We showed that both in simulated data and in an NHP neural dataset, the MED was able to simultaneously detect and classify eye movements in a way that successfully combined information across data modalities.

### 4.1. Bi-Directional Performance Improvement

A successful multimodal decoder should take advantage of the information present in all available data modalities. To show that the MED does this, we performed an extensive performance analysis in which channels of both modalities were incrementally included as input to the MED. On both simulated and real NHP data, this analysis showed that MED performance improves bidirectionally – both when LFP channels are added to a fixed number of spike channels, and when spike channels are added to a fixed number of LFP channels. In the NHP dataset, although spike-only performance was generally better than LFP-only performance, multimodal performance was still significantly better than both. These results suggest that LFPs contain information about both event timing and event class that is not present in spikes, and vice versa.

### 4.2. Cross-Modal Scaling

As mentioned in section 2.2, we modified the maximum-likelihood estimator of event times and classes by adding a cross-scale combination parameter. As we explained, this is necessary when we have a combination of discrete and continuous modalities, because the likelihood in the former provides a probability measure while the likelihood in the latter requires integration over a range of values to give a probability. Thus combining likelihoods of discrete and continuous modalities is sensitive to scalings of the latter. Indeed, scaling a Gaussian signal will in turn scale its likelihood, even though its signal-to-noise ratio (SNR) is unchanged.

To see why this is, consider a signal *y* = *s* + *n*, where *s* = *b* is a constant signal and *n* ∼ N (0, *σ*^2^) is noise. The SNR of this signal is *b*^2^*/σ*^2^, and the likelihood of observing *y* = *b* is given by the peak of the Gaussian PDF 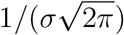. If *b* = 1 and *σ*^2^ = 1, then the SNR of *y* is 1 and the likelihood of observing *y* = 1 is 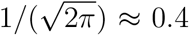. If we then multiply *y* by 2, this is equivalent to setting *b* = 2 and *σ*^2^ = 4. The SNR is still 1, but the likelihood of observing *y* = 2 is now 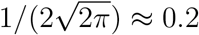 So, even though scaling *y* does not change its SNR, it *does* change its likelihood. This is problematic because ideally the contribution of Gaussian channels to the MED output should depend only on their SNR, and not on their scale. In practice, we found that a single parameter weighing the contribution of all Gaussian channels together was sufficient to properly combine their information with the point-process channels, thereby alleviating this issue.

In addition to the above mathematical reason, another practical reason for using this scaling parameter is model mismatch in real neural datasets. If any of the model parameters are incorrectly estimated in a systematic way across a single modality, then the learned model will not properly weigh the contributions of each modality. For example, it is possible that there are events unrelated to eye movements that also elicit spatiotemporal responses in both types of channels. For Gaussian channels, this can be accounted for by the noise variance parameter *σ*, but Poisson models do not have a way to account for this kind of variance. This phenomenon could cause one scale to be ignored even if it has information to contribute.

### 4.3. Applications and Future Directions

The application of the MED in a real-time BCI is an important research direction. While motor BCIs to date have focused on continuous movement decoding, the development of cognitive BCIs will in many contexts require event detection in order to perform decoding. For example, in the context of a stimulus-based decision task, one would first need to detect a stimulus onset in order to then decode decision-related information, such as the decision itself or the associated confidence [64–67]. Thus, the MED can enable multimodal cognitive BCIs for decision making in real-world applications where event times are unknown. The MED can also help extend multimodal motor BCIs to naturalistic setups where task-related events, such as movement onset, must be detected before decoding is possible. In addition to spiking and LFP, the application of MED to other neural modalities such as intracranial EEG or electrocorticography, which have high potential for translation, is also an important future direction [22; 68–82].

In section 2.2, we assumed conditional independence between the spiking and LFP modalities, conditioned on event times and classes, in order to make the derivations tractable. Since the MED successfully combined information across data modalities in both event detection and classification, we can conclude that this is a reasonable assumption. The fact that this conditional independence assumption between LFPs and spikes can be reasonable has also been shown in various prior studies (e.g., [31; 35]). Further, without this assumption, the number of parameters can become prohibitively large if interdependencies between every channel pair are considered, thus raising the possibility of overfitting to training data – especially in neural datasets that can be limited in sample size. Nevertheless, future work can consider modeling such interdependencies while regularizing or sparsifying them to avoid overfitting.

## Appendix A. Model Parameter Estimation

In order to make use of the MED, the model parameters of each channel must be estimated. Prior work has developed a way to estimate spatial and temporal parameters for Poisson GLMs [60]. This method can also be used with Gaussian linear models. Building on this prior work [60] for Poisson GLMs, we estimate the parameters in our multimodal model as follows.

The parameters that we must estimate are the temporal responses *r*(*t*), the spatial responses *ϕ*, the Poisson baseline parameters *α*_*i*_, and the Gaussian noise variances *σ*_*j*_. We can estimate these parameters by maximizing the likelihood of a training dataset over these parameters. For Poisson signals, this can be done efficiently via iteratively re-weighted least squares, and for Gaussian signals, this can be done via simple linear regression. However, these methods require that our models are log-linear in the parameters for Poisson channels, and linear in the parameters for Gaussian channels. Consequently, we must rewrite our multimodal model in a way that is suitable for parameter estimation.

In our models (1) and (3) the temporal responses *r*(*t*) are not in a parametric form. We can address this by parameterizing them as a weighted sum of *B* basis functions:

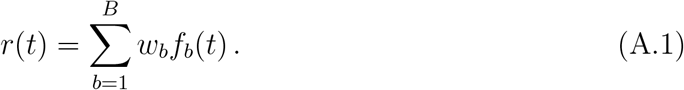

In this way, the temporal response is parameterized by the weights *w*_1_, …, *w*_*B*_. In this work, the basis functions *f*_*b*_(*t*) are 10 truncated Gaussians with a standard deviation of 100ms, with means evenly spread out from -500ms to 500ms relative to the event time *t*_*s*_.

Now that our model has been parameterized, we need a way to represent the independent variables, the event classes *s* and times *t*_*s*_, in a way that is suitable for generalized linear regression. To do this, we define a signal *d*_*n*_(*t*) that encodes both the event classes and the event times:

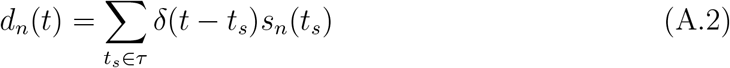

Here, *s*_*n*_(*t*) is the *n*th element of the one-hot vector *s* at time *t* and *τ* is the set of event times. Using the parameterization of *r*(*t*) in (A.1) and the event-encoding signal *d*(*t*) in (A.2), the terms 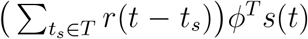 in (5) can be rewritten as a matrix multiplication, in a manner similar to [60]:

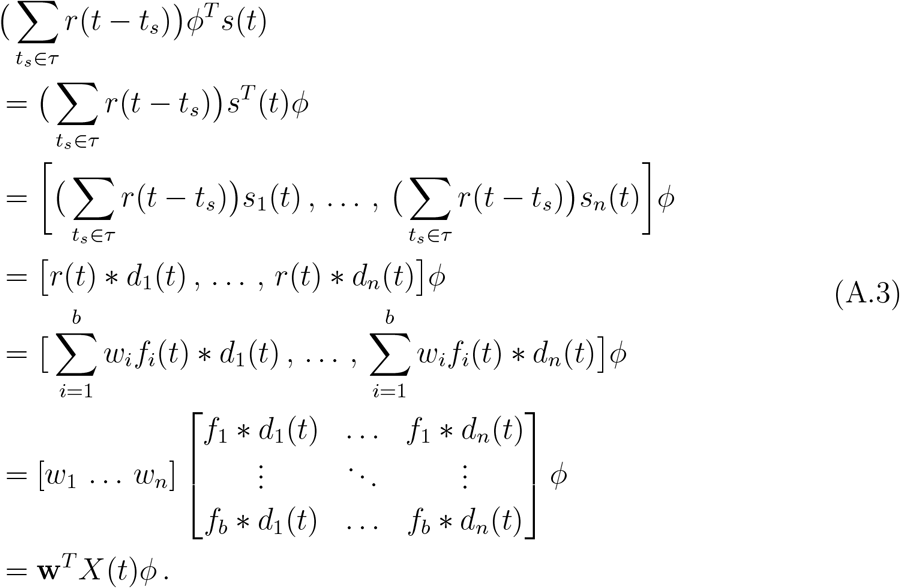

Here, ∗ is the convolution operation and **w** = [*w*_1_ … *w*_*b*_]^*T*^ is a vector of the temporal response parameters. The matrix *X*(*t*) encodes the independent variables *t*_*s*_ and *s*, which will be known for training data, along with our chosen basis functions *f*_*b*_(*t*). Substituting this into (1) and (3), we obtain a generalized bilinear model for point process channels:

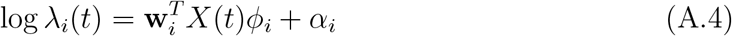

and a bilinear model for Gaussian channels:

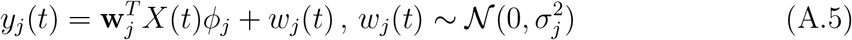

**w** and *ϕ* are parameter vectors that represent the temporal and spatial responses respectively, and must be estimated from the training dataset. Generalized bilinear models have previously been used to model the spatiotemporal responses of spiking neurons [60; 83].

If we fix the value of *w*, then we can learn *ϕ* by fitting a linear model or Poisson GLM with 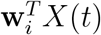 *X*(*t*) as the independent variable and the training data, either Gaussian or point-process, as the dependent variable. We can similarly learn *w* by fixing *ϕ* and using *X*(*t*)*ϕ*_*i*_ as the independent variable. In this work, we fit parameters by iteratively fixing either the spatial or temporal response and fitting the other, as in prior works [60; 83]. Parameters were initialized to vectors of all ones.

For point-process signals, *α* is learned as the bias parameter in the GLM, and for Gaussian signals, *σ* is the standard deviation of the residuals after the parameters are learned.

